# A recipe for recovery: Diet as a decisive tool in species conservation

**DOI:** 10.1101/2024.12.05.625804

**Authors:** Noa Lavie, Yael Levinson, Jessica M. Rothman, Dror Hawlena

**Affiliations:** Risk-Management Ecology Lab, Department of Ecology, Evolution & Behavior, The Alexander Silberman Institute of Life Sciences, The Hebrew University of Jerusalem.; Department of Anthropology, Hunter College of the City University of New York, New York, New York, USA; Science Division, Israel Nature and Parks Authority, Jerusalem, Israel

**Keywords:** Nutritional ecology, *Vachellia tortilis*, Ungulate reintroduction, Endangered species conservation, Right-angled Mixture Triangle, *Gazella arabica acacia*

## Abstract

Animals consume foods in specific quantities and ratios to meet a multi-dimensional nutrient target that maximizes their fitness. Attempts to reach this target in an ever-changing nutritional landscape often influence animal foraging behavior, habitat choice, and food-web interactions. Consequently, understanding the nutritional ecology of a species is instrumental in developing efficient management plans for its conservation. We quantified the dietary considerations of a critically endangered population of Acacia gazelles (*Gazella arabica acacia*), and compared their diet and nutrition to that of the sympatric *Gazella dorcas,* using behavioral observations coupled with nutritional and secondary metabolite quantification, and morphological measurements of trees. Acacia gazelles mainly consumed resources from two subspecies of the umbrella-thorn acacia (*Vachellia tortilis*) that differ in morphology, nutritional composition, and phenology, and preferred browsing on specific trees based on their nutritional and defensive traits. Gazelles maintained a narrow nutritional target, tightly regulated their protein intake, and aimed to increase consumption of non-structural carbohydrates. Dorcas gazelles consumed a very different diet from the Acacia gazelle, but converged to a similar intake target. By considering both dietary and nutritional data, we conclude that suitable habitats for Acacia gazelle reintroduction must include sympatric populations of the two *V. tortilis* subspecies, that nutritional deficiency is unlikely to play a major role in explaining the slow rate of population recovery, and that the seemingly low competition between the two gazelle species does not necessitate controlling the Dorcas gazelle population. Our results link species-specific dietary strategies with resource selection and resulting interspecific interactions, demonstrating the importance of nutritional ecology as a major conservation tool.

## 1. Introduction

Wildlife feed on various foods in specific ratios and quantities to meet a multifaceted nutrient “intake target” that maximizes their lifetime fitness (sensu; Simpson and Raubenheimer 2012) and may vary with age, sex, and physiological state (e.g., Simpson and Raubenheimer 1997, Raubenheimer and Simpson 1999, Hawlena and Schmitz 2010). Attempts to meet this target in an ever-changing “nutrient landscape” (i.e., the spatial and temporal distribution of nutrients available to animals within their environment; Youngentob et al. 2026) may influence animals’ spatial activity and behavior (Bazazi et al. 2010, Johnson et al. 2017), and affect their interactions with competitors, natural enemies, and anthropogenic risks (Kay et al. 2010, Coogan and Raubenheimer 2016, Machovsky-Capuska et al. 2019, Nie et al. 2019, Gazzola and Balestrieri 2020). Moreover, human activities may alter the abundance, quality, and distribution of food resources (Acevedo-Whitehouse and Duffus 2009, Hamann et al. 2021, Fehlmann et al. 2021), possibly limiting wild animals from meeting their intake target (Armstrong and Perrott 2000, López-Bao et al. 2010). Thus, uncovering the dietary and foraging considerations of endangered species in the wild is key to address the challenge of conserving them (Birnie-Gauvin et al. 2017).

We adopted this dietary-centric approach to assist in conserving a critically endangered Acacia gazelle (*Gazella arabica acacia*) population in the Southern Arava, Israel. Acacia gazelles are a genetically unique and geographically isolated population of Arabian gazelles (*Gazella arabica*; Hadas et al., 2015; Fig. 1). This remnant population of about 40 individuals is restricted to two small floodplains, and struggles to recover despite substantial conservation efforts (Shalmon 1991, Blank 1996, 2000, Polak et al. 2023). Historical references suggest a much larger distribution before the 1950s, when the entire population was almost extirpated by hunting (Mendelssohn 1974). However, not enough is known about this historic distribution to predict the gazelles’ habitat requirements and locate suitable reintroduction sites. In 2006, the last 13 individuals were enclosed within a predator-proof area, alongside 82 Dorcas gazelles (*Gazella dorcas*) incidentally trapped during the installation. It was hypothesized that nutritional limitations, due to intensifying competition with the cohabiting Dorcas gazelle and food shortage, might impede the population’s recovery (Shalmon et al. 2020, Breslau et al. 2020). Thus, revealing the Acacia gazelle’s natural diet and nutritional needs, and comparing them to those of the Dorcas gazelle, may allow the development of an effective plan to save this unique population.

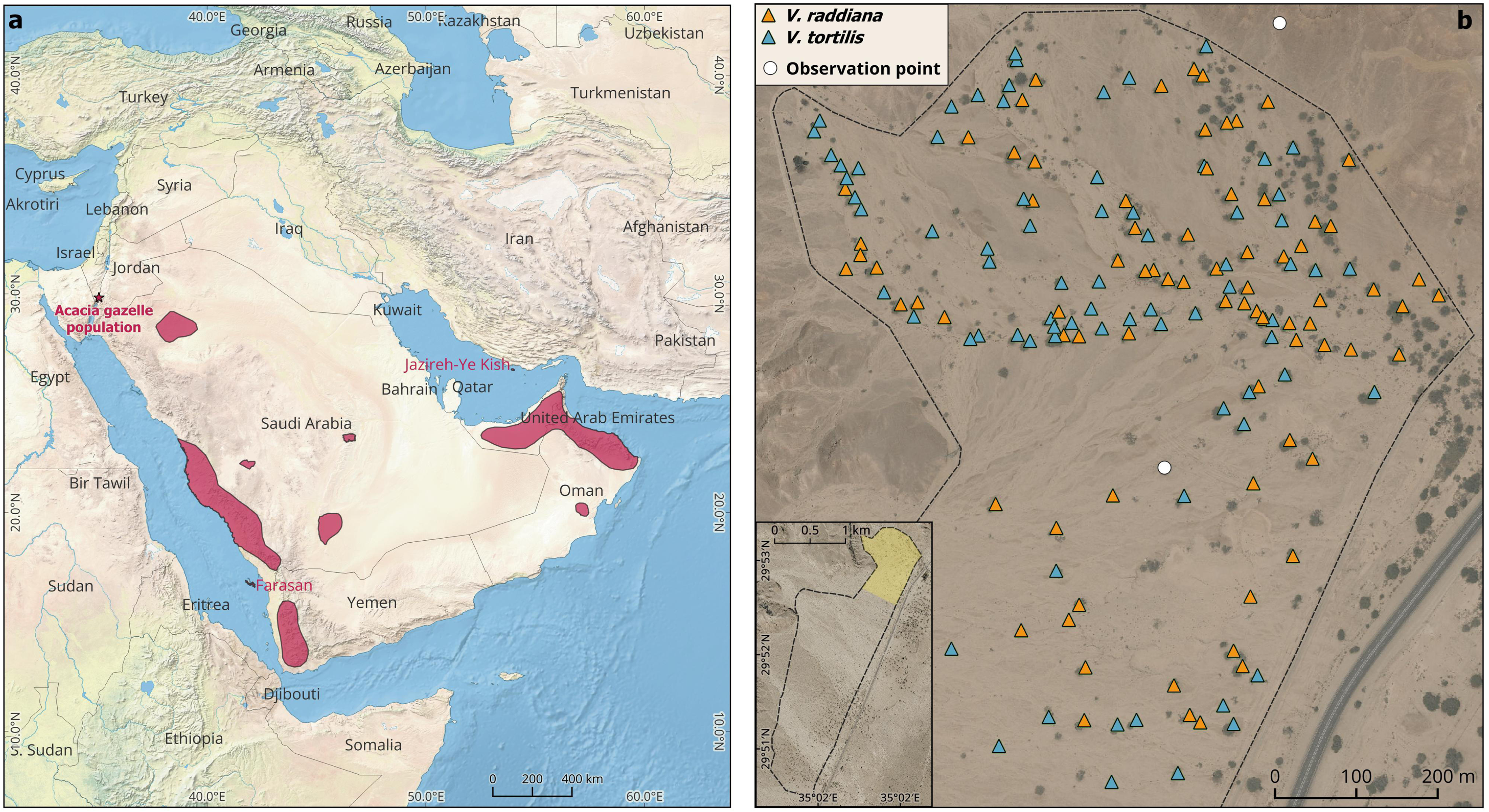

Attempts to quantify nutritional needs and other dietary considerations of wildlife in the field traditionally focused on a single currency such as biomass, energy, or protein consumed (Schoener 1971, Charnov 1976, White 1993). However, natural foods are complex mixtures of macronutrients and micronutrients in varying levels of availability and digestibility. Animals often need to ingest several food types, and occasionally digestive assistant agents (i.e., substances or organisms that facilitate the digestion of other foods) to meet their multidimensional nutritional target (Zaguri et al. 2024). Thus, focusing on a single food currency may provide only a limited understanding of the animals’ true dietary considerations (Simpson et al. 2004, Felton et al. 2009a, 2018). Progress in nutritional ecology, especially the development of nutritional geometry and the right-angled mixture triangle (RMT) method, which uses proportions rather than amounts of foods consumed to quantify wildlife intake targets, marked a breakthrough in our ability to use nutritional ecology to inform conservation (Raubenheimer 2011, Raubenheimer et al. 2015). Yet despite the clear potential, RMT has been used almost exclusively in primate research (Raubenheimer et al. 2015, Johnson et al. 2017, Takahashi et al. 2019, DePasquale et al. 2023), and only rarely to quantify intake targets of endangered wildlife (e.g., Raubenheimer and Simpson 2006, Nie et al. 2015, Panthi et al. 2019, Hecker et al. 2021), or ungulates (Raubenheimer et al. 2014, Spitzer et al. 2023). In these rare cases, the limited ability to directly observe the animal’s feeding behavior and to quantify the nutritional composition of foods has led to the development of proxy methods like fecal (Nie et al. 2015, Hecker et al. 2021, Spitzer et al. 2023) or rumen and gut content analyses (Felton et al. 2021) to quantify the diet, and the use of crude nutritional estimates at the species level to assess the food nutritional content (Costello et al. 2016, Shrestha et al. 2020, Spitzer et al. 2023, 2024). These proxy methods are informative but ignore temporal and intra-specific variation in the availability and nutritional content of food resources, leading to cruder intake target estimates that are less applicable for management purposes.

Focusing solely on the nutritional composition without considering other food characteristics may produce an incomplete understanding of animals’ dietary considerations. Plants are often defended by various chemical (e.g., tannins; alkaloids; salts) and physical defenses (e.g., thorns; opaline phytoliths; trichomes), and include components of little nutritional value (e.g., lignin, Barboza et al. 2009). These traits may alter herbivores’ ability to ingest and digest foods, possibly influencing the herbivore’s dietary choices (Freeland and Janzen 1974, Cooper and Owen-Smith 1986). Consequently, attempts to uncover the dietary strategies of endangered ungulates must include quantification of nonnutritive plant traits that influence food choice.

Our objectives were to determine the natural diet and nutritional requirements of the Acacia gazelle and to compare them to those of the Dorcas gazelle. We accounted for the methodological considerations previously described by observing the Acacia gazelles’ foraging behavior in the field, collecting samples from all plants consumed and from similar reference resources that were not consumed, and measuring the morphological traits and tannin composition of *Vachellia* trees. We characterized the chemical composition of all samples using wet chemistry and Near Infra-Red Spectroscopy (NIRS), and employed the RMT method to estimate the gazelle nutritional intake target. Ultimately, the resulting insights provide a rigorous empirical basis for revising the conservation management plan for the critically endangered Acacia gazelle.

## 2. Materials and Methods

### 2.1 Study site and gazelle populations

The study was conducted within the Acacia gazelle predator-proofed enclosure in Yotvata Nature Reserve, Israel (29°51’-29°53’N, 35°01’-35°03’E; Fig. 1). This hyper-arid site is located within the Jordan Rift Valley about 40 km north of the Gulf of Aqaba. The mean annual temperature is 24°C, and the average annual precipitation is 40.5 mm with big fluctuations between years. All precipitation occurs between October to May. The site includes two alluvial fans dominated by scattered *Vachellia* (formerly *Acacia*) trees and their common semi-parasite *Loranthus acaciae*. Several halophyte shrubs, mainly *Haloxylon salicornicum*, *Lycium shawii*, *Salsola imbricata,* and *Nitraria retusa*, dominate the exposed stony gaps between the *Vachellia* stands (Baharav 1982). Two *Vachellia tortilis* subspecies coexist in this site *(V. tortilis tortilis* and *V. tortilis raddiana).* Locally, the two subspecies exhibit consistent differences in their chemical and morphological traits, and habitat requirements (Halevy and Orshan 1972, Rohner and Ward 1997). For example, *V. raddiana* typically reaches a height of 4-7 m, while *V. tortilis* is a shorter tree reaching a height of only 2-3 m (Ward and Rohner 1997). Moreover, the two trees have very distinct phenology, producing flowers and fruits at different times (Halevy and Orshan 1973). Thus, these two subspecies are treated locally as distinct taxa (Zohary 1972). Following this approach, we treat the two *Vachellia* trees as distinct resources, hereafter referred to as *V. tortilis* and *V. raddiana*.

An Acacia gazelle population of 35 individuals was first discovered at our study site in 1964 (Mendelssohn 1974, Yom-Tov and Ilani 1987, Mendelssohn et al. 1997). This population was discovered only after all other Acacia gazelle populations in Israel were extirpated. Since then, the Israel Nature and Parks Authority (INPA) has monitored and attempted to restore the population using various management interventions like supplementary feeding with natural or artificial foods, predator culling, and removal of calves in an attempt to establish a breeding colony in captivity (Blank 2000). Despite those substantial efforts, the population failed to recover (Polak et al. 2023). In 2006, after a dramatic population decline, the INPA fenced the remaining 13 gazelles together with 82 Dorcas gazelles (*Gazella dorcas*) incidentally trapped during the installation in a 3.5 km^2^ predator-proofed enclosure (Polak et al. 2023). By 2013, both Acacia and Dorcas gazelle populations reached a maximum density of 42 and 173 individuals, respectively. However, in the winter of this year, the fence was breached by flood water, enabling wolves to penetrate and kill about 30 Acacia gazelles and 65 Dorcas gazelles (Polak et al. 2023). During our field observations in the summer of 2014, 27 Acacia gazelles and 138 Dorcas gazelles cohabited the enclosure. Between 2015 to 2019, the INPA actively removed 161 Dorcas gazelles from the enclosure as a precautionary step against interspecific competition, leaving only 36 Dorcas gazelle individuals in the enclosure (Polak et al. 2023). During our field observations in 2021-2022, only about 30 Acacia gazelle and about 40 Dorcas gazelle cohabited the enclosure (Polak et al. 2023).

### 2.2 Gazelle feeding observations and forage sampling

We conducted group observations of the gazelles’ feeding behavior in monthly intervals during one year (2021-2022). We observed the gazelles from two hides (Fig. 1b) during their peak foraging activity in the early morning and late afternoon for a minimum of 3 consecutive days each month. Each observation session lasted for 128±42 minutes. We recorded the number, age, and sex of feeding gazelles, the feeding time, and whether gazelles were feeding from the ground or from the canopy, along with the exact location and type of each food resource (see Appendix 1 for details). After each observation, we collected samples from all plant resources that were consumed by the gazelles and immediately transferred the samples to our field lab. There, we weighed and oven-dried the samples (60°C, 48 h), and re-weighed them for dry mass estimations. We milled all samples through a 1 mm sieve using a laboratory grinding mill (Kinematica POLYMIX PX-MFC 90D), and stored the milled samples in the freezer until further analyses.

To compare the diet of Acacia gazelles to that of Dorcas gazelles, we used continuous time-budget observations on focal individuals of the two gazelle species. In each observation, we tracked a focal individual for 15±9 minutes and then switched to another gazelle. We collected these data in the summer of 2014 (Aug–Oct), prior to the removal of the Dorcas gazelles, using the same times of day and observation hides as in the 2021–2022 period. (see Appendix 1 for details). By comparing the feeding behavior and calculated intake targets of Acacia gazelles from August–October 2014 with those from the same months in 2021–2022, we were able to evaluate the two observation methods and assess whether nutrient intake was affected by fluctuations in the number of cohabiting Dorcas gazelles.

### 2.3 Chemical analyses

We used classic chemical quantification methods to estimate the nutritional content of 209 out of 400 collected food samples (Table S1). Each of the 209 samples was analyzed for total non-structural carbohydrates (NSC), crude protein, lipids, fiber, ash, and tannin content. NSC were quantified using the phenol-sulfuric assay with acid extraction (Dubois et al. 1956, Landhäusser et al. 2018). For crude protein estimation, we estimated the Nitrogen (N) content by combustion, using a Leco FP-528 Nitrogen Analyzer (Michigan, USA), and multiplied the N content by the widely used 6.25 conversion factor (Jones 1931, Rothman et al. 2012). Lipids were measured using ether extract in an ANKOM Fat Analyzer (New York, USA). Fiber fractions were determined using the sequential detergent fiber assay (Van Soest et al. 1991) in an ANKOM A200 Fiber Analyzer: Neutral Detergent Fiber (NDF), which contains non-soluble fiber (hemicellulose, cellulose, and lignin), Acid Detergent Fiber (ADF, cellulose and lignin), and Acid Detergent Lignin (ADL). Samples were burned at 500–550°C for ash measurement (Rothman et al. 2012). Finally, the relative content of condensed tannins was estimated using the acid-butanol assay after extracting samples with 70% acetone (Porter et al. 1985, Hagerman and Butler 1994, Rothman et al. 2009). Additionally, we chose 56 samples of the food resources observed in 2014 (8-10 samples of each food resource) for mineral quantification to assess the ash composition. Minerals were assessed via atomic absorption spectroscopy after microwave digestion at Dairy One Commercial Forage Laboratory (Ithaca, NY, USA). Nutrient estimations are presented on a 105°C dry matter basis. We estimated the available protein in a subset of samples of these main resources to rule out possible digestibility biases of our crude protein measure due to protein binding in fiber. Therefore, we measured N content in the ADF residue, multiplied by the same factor, and subtracted from the crude protein (Licitra et al. 1996, Rothman et al. 2008), see Fig. S1 for details.

We constructed NIRS models for *Vachellia* leaves and pods based on 151 samples analyzed by wet chemistry (Table S1). These samples were chosen based on quantity and Near Infrared Spectroscopy (NIRS) spectral variation to minimize misspecification of the calibration equations. *Vachellia* flowers, *Loranthus* leaves and flowers, and perennial shrubs were analyzed only by wet chemistry due to the low number of samples, and the average nutrient content was applied. For NIRS spectra collection, we dried all samples for 48h. After homogenizing the sample, we transferred a ∼50 mg subsample into a 1 ml clear glass shell vial and scanned each sample using an Antaris II FT-NIR analyzer. We constructed two NIRS analyses using TQ analyst software. The first analysis included models for the absolute concentrations of all measured nutritional components (Protein, Lipids, NSC, NDF, ADF, and ADL). The second analysis was for the relative concentration of tannins. NIRS models’ protocols and performances are presented in Appendix 2.

### 2.4 Physical measurements of *Vachellia* trees

We randomly chose 20 *V. tortilis* and 20 *V. raddiana* trees in the northern part of the enclosure. Half were observed browsed by Acacia gazelles during the winter, when mainly foliage is available (hereafter “browsed”), and half were randomly chosen reference trees from all trees for which we did not detect feeding in the previous year (“unbrowsed”). For each tree, we divided the canopy into 4 equal-sized sections and measured 2 randomly chosen branches in each section. All sampled branches were within the feeding range of an adult Acacia gazelle (up to 1.7 m, Fig. S2). Following the protocol described by Rohner & Ward (1997), on each branch, we counted all leaves, and all thorns longer than 1mm (perpendicular to the branch) within 50 cm from the branch’s distal end. We classified thorns as either straight (spines) or curved (prickles), and the lengths of all thorns >5mm were measured perpendicular to the branch. A sample of the 5 biggest leaves was measured for length and width as well.

### 2.5 Data analysis

We estimated the Acacia gazelles’ nutritional target and resources’ composition using a RMT (Raubenheimer 2011), of the relative contents of the three main macro-nutrients in the gazelle diet – Protein, NSC, and digestible fiber (hemicellulose and cellulose). We calculated the daily nutritional target by multiplying each observation unit’s (plant) nutritional values by the time and number of gazelles spent feeding from it, divided by the total observed daily feeding time. To unravel whether gazelles were selecting for specific macro-nutrients, we conducted a permutation test for the gazelles’ intakes. We summarized each food resource’s mean nutritional concentrations and its total feeding time within each observation session. Then, we permuted these feeding times 1000 times and recalculated the permuted intake target to create a null intake distribution. We calculated the P-value as the fraction of permuted values exceeding the observed intake, and report the median across 1000 permutation runs. We consider this approach conservative, yet realistic, as we have only permuted consumption times of resources that were selected by the gazelles to be included in their diet, depicting the species diet breadth shaped by its evolutionary history. Therefore, we suggest that marginal p-values are meaningful. To calculate the diet and intake target of the two gazelle species as observed in 2014, we calculated the gazelles’ relative use of each resource for the whole period and applied the mean nutritional values of this resource for the respective period.

We performed a Kruskal-Wallis test followed by a Bonferroni-corrected Conover-Iman post-hoc test (Conover and Iman 1979) to compare the means of all macro-nutrients, minerals, and tannin contents of the most prominent resources in the diets of both gazelle species as observed in 2014 (i.e., *Vachellia* leaves and pods, *Haloxylon salicornicum*, and *Loranthus acacia* leaves). We calculated fiber nutritional values (i.e., hemicellulose, cellulose, and lignin) by subtracting the chemical sequential fractions (NDF-ADF, ADF-ADL, ADL, respectively). Since subtraction introduces a greater potential error, we also report the raw values in the supplementary, where applicable.

We used PCA to evaluate inter-specific variance between the *Vachellia* taxa. To uncover differences between the 20 browsed and 20 unbrowsed trees of each taxon, we applied a separate PLS Discriminant Analysis (PLS-DA, Barker & Rayens 2003) within each *Vachellia* subspecies, since the PCA revealed a large inter-specific variability. We standardized the data before both ordinations. For the morphological traits measured, we averaged all traits for all branches of each tree. We analyzed the thorns data using counts within two length categories (with a 5 mm cutoff), resulting in four thorn categories (small spines, big spines, small prickles, and big prickles). Nutritional data were obtained for each tree from the NIRS model predictive results, except for two of the *V. raddiana* trees that had only morphological measurements and lacked nutritional data. Consequently, we removed these two *V. raddiana* trees from this analysis.

To assess the difference between browsed and unbrowsed trees in the PLS-DA analysis, we performed a permutation test on these browsing categories. We tested for significant cluster separation using the Mahalanobis distance between the two categories’ centroids based on the pooled covariance matrix. We also tested for significance the proportion of explained variance between the browsing categories (*R*^2^) and *DR*^2^, correcting for false-negatives resulting from out-of-bounds estimation (Westerhuis et al. 2008). To uncover which variables are the most influential in creating these patterns, we estimated each variable’s importance by randomly permuting only its nutritional values and conducting the same analyses. We compared the means of permutations’ *R^2^*, *DR^2^*, Mahalanobis distance between clusters, and the fraction of significant permutations for cluster separation.

All statistical analyses were conducted in R 4.3.0 software (R Core Team 2023). Kruskal-Wallis and Conover’s tests were performed using ’kruskalTest’ and ’kwAllPairsConoverTest’ from ’PMCMRplus’ package. PCA was fitted using ’prcomp’ from ’stats’ and PLS-DA was fitted using ’plsda’ from the ’mixOmics’ package (Rohart et al. 2017).

## 3. Results

Six different plant species were consumed by Acacia gazelles during the study. Their diet consisted mainly of different resources from two *Vachellia* subspecies, which together constituted about 88% of the annual gazelle diet, and no less than 70% of the monthly diet (Fig. 2a). The consumed resources differed substantially in their nutritional and mineral content (Kruskal-Wallis *p*<0.001 for all, see tables S3 and S4 for post-hoc comparisons).

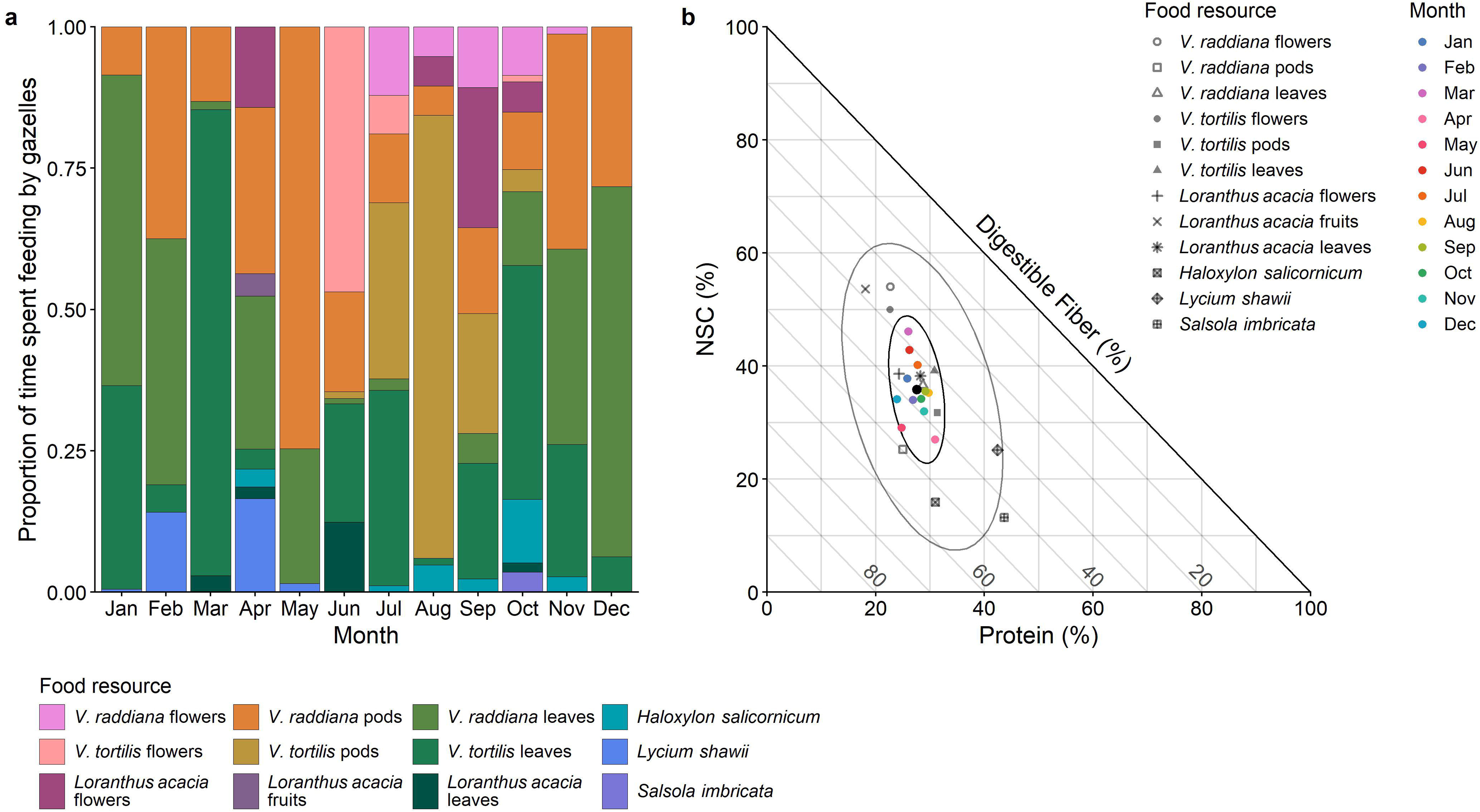

The two *Vachellia* subspecies differed substantially in their chemical and physical characteristics. *V. raddiana* trees had bigger leaves, while *V. tortilis* trees had higher densities of prickles, and higher concentrations of proteins and lipids in their leaves (Fig. S3). PCA results revealed high intra-specific variability in spine counts, and the contents of NSC, fibers, and tannin in the leaves of both subspecies (Fig. S3).

*Vachellia* subspecies differed also in their phenology. *V. tortilis* trees flowered while shedding their leaves in early summer, whereas *V. raddiana* trees started flowering in late summer and throughout the fall without completely shedding their leaves. *V. tortilis* pods developed immediately after the blooming period and ripened in late summer, while *V. raddiana* green pods remained on the branches during winter and ripened only in the next spring (Fig. 3c). *V. tortilis* produced flowers and fruits in shorter and more synchronized time windows than *V. raddiana* (Fig. 3c). The rate of gazelles’ visits to the *Vachellia* trees, and their relative feeding time were tightly associated with the above-listed phenological differences (Fig. 3). For instance, gazelles seemed to prioritize feeding on flowers when those were available. During the peak *V. tortilis* bloom in June, gazelles shifted their diet toward *V. tortilis* flowers, which made up to 47% of their diet, despite an abundance of green *V. raddiana* pods (Fig. 3).

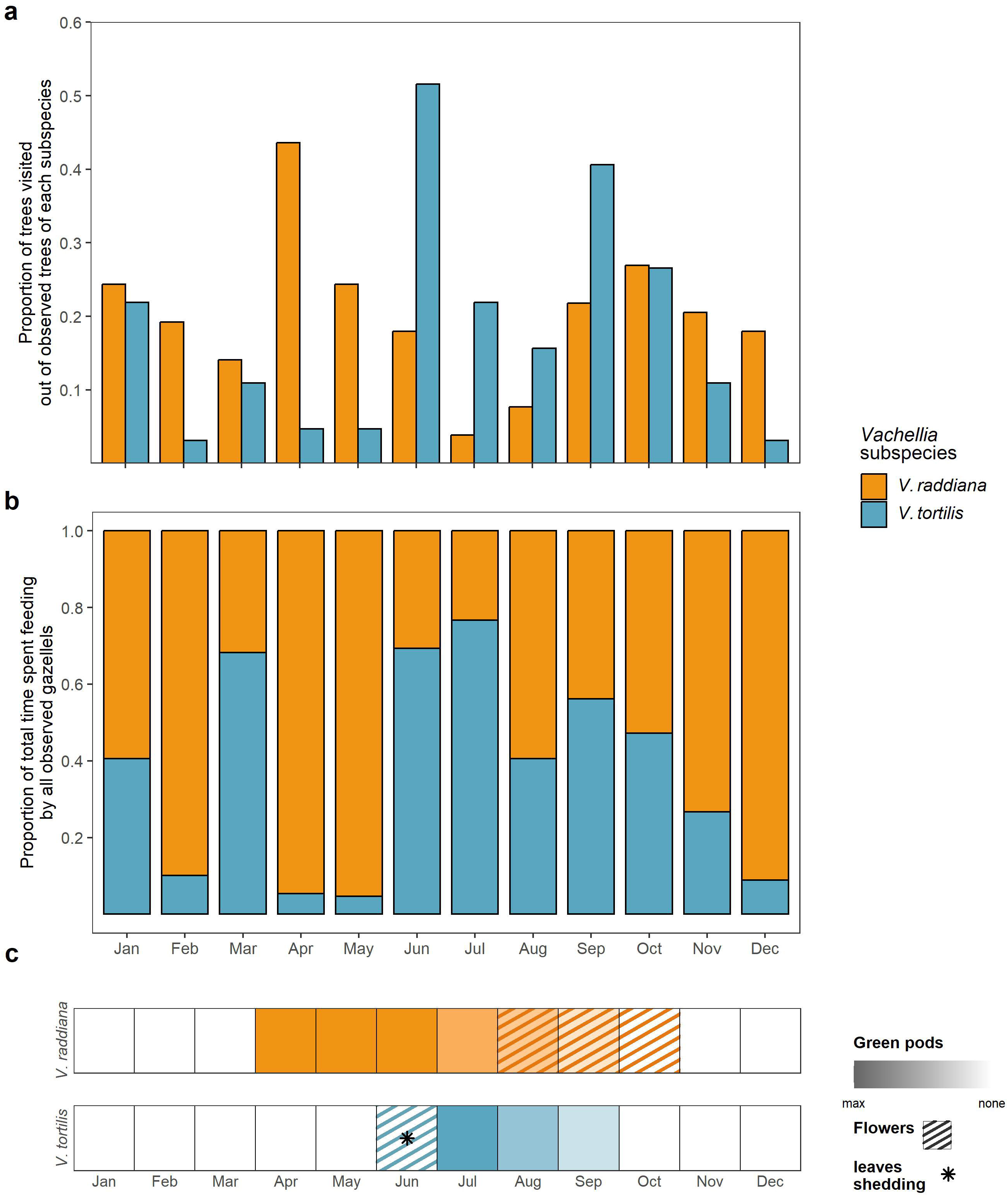

The variation in the gazelle’s monthly nutritional intake was much smaller than the variation in the nutritional landscape (Fig. 2b, Table S3). Gazelles achieved this tighter balance by eating differentially from the different available food resources. For example, in February, the gazelles ate *V. raddiana* leaves from the canopy and pods from the ground, as well as *L. shawii* leaves in order to reach their nutritional target (Fig. 2, Fig. S4). This tight intake target was reached despite substantial differences in resource availability (Fig. 3, Fig. 2). From all macro-nutrients measured, protein intake varied the least (Table S3). The permutation test shows no significant selection for protein-rich foods (*p*=0.47). Thus, the precise protein intake may reflect the smaller variation in protein compared to other macro-nutrients in the consumed resources (Fig. S5). Similarly, we observed no selective feeding for digestible fiber (*p*=0.33, Fig. S5). However, the permutations test revealed that gazelles tend to select for non-structural carbohydrates, as the observed intake was higher than the expected intake by random feeding (*p*=0.07). During summer, gazelles increased their relative NSC intake by prioritizing *V. tortilis* NSC-rich flowers (Fig. 2). Still, gazelles kept their tight protein intake by consuming *V. raddiana* pods and leaves from both *Vachellia* subspecies.

We observed variation in the use of individual *Vachellia* trees, of both subspecies, where some trees were heavily browsed while others were ignored (Fig. S6). The browsed and un-browsed *V. tortilis* trees differed substantially (Fig. 4a, *M_d_*=3.28, *p*=0.014). The variation in the browsing category explained by the model was 75%-78% and was significantly higher than random for both fit-measures (*R^2^* =0.75, *DR^2^* =0.78, *p*=0.021, 0.019, respectively). The separation was mainly along the first ordination axis (LV1) driven predominantly by the high fiber fractions and bigger leaf size (expressed as width and length measures) of the un-browsed trees, and the higher protein as well as higher number of leaves of the browsed trees (Fig. 4a,c). Browsed trees were also characterized by having a lower density of big prickles, and un-browsed trees had a relatively higher density of small prickles. Hemicellulose, big prickles, and protein were the variables for which randomization had a major effect on the model performance and cluster separation (Table S5). We did not find significant differences between the browsed and un-browsed *V. raddiana* trees (Fig. 4b). Using raw analytical fiber fractions instead of the nutritionally calculated fractions slightly decreased the performances of both models, but it remained significant for *V. tortilis* (Fig. S7).

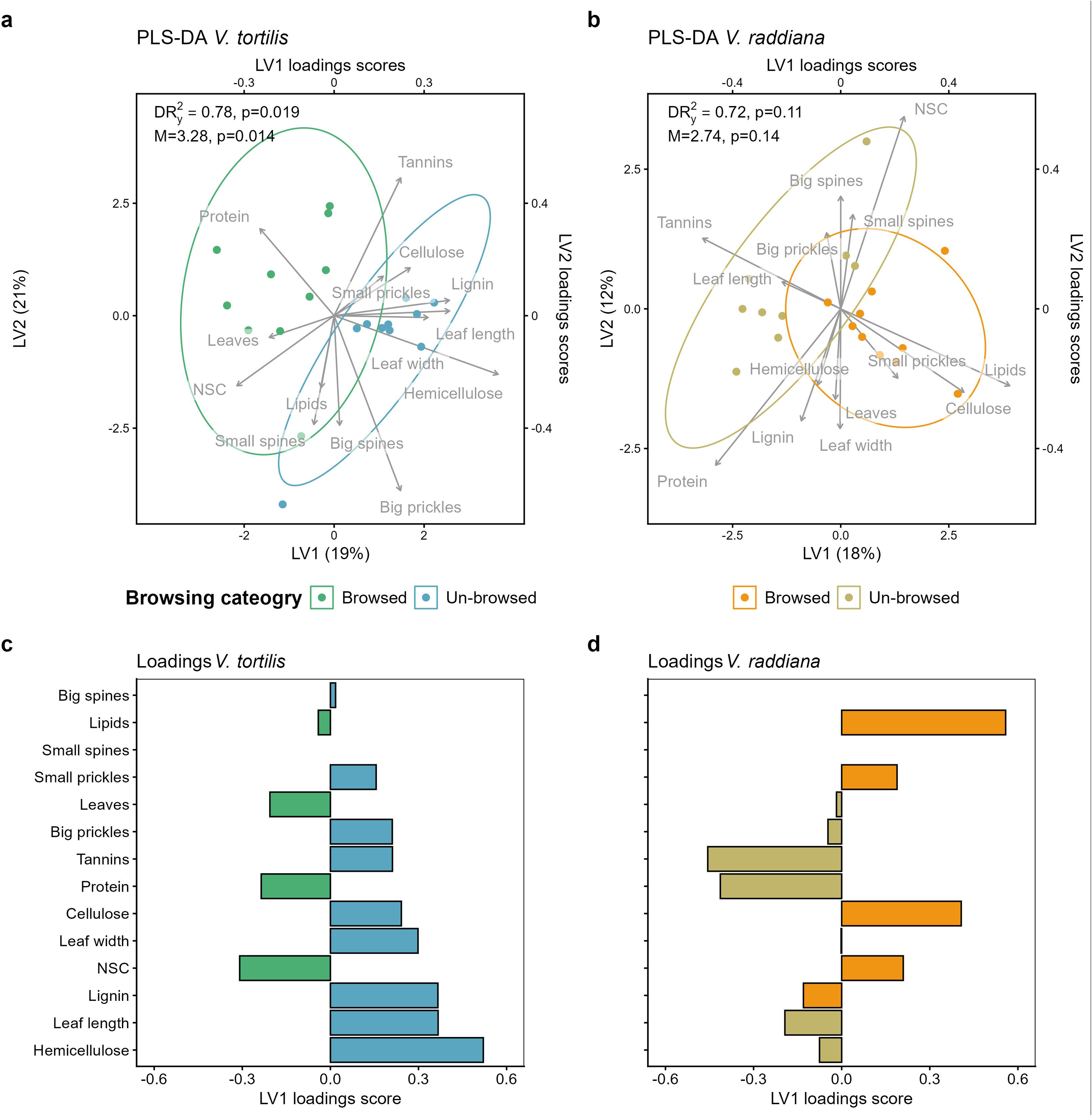

We found major differences between the diet compositions of the two gazelle species. Acacia gazelle foraged predominantly on two *Vachellia* subspecies, while Dorcas gazelles’ diet included more shrubs, especially *Haloxylon salicornicum*. When feeding from the two *Vachellia* trees, Acacia gazelles consumed a much higher proportion of *V. tortilis* in comparison to Dorcas gazelles, which fed almost exclusively from *V. raddiana* (Fig. 5a). Acacia gazelles fed more on the foliage from both *Vachellia*s, while Dorcas gazelles fed predominantly on *V. raddiana* pods and flowers from the ground. Additionally, Acacia gazelle fed on *Loranthus acacia* while Dorcas gazelle did not use this resource (Fig. 5a). Despite the large dietary differences, the two gazelle species converged to a similar intake target. Both gazelle species achieved very similar protein intake, but Acacia gazelles reached higher NSC and lower digestible fiber than Dorcas gazelles (Fig. 5b). Our two independent measurements of Acacia gazelle intake targets in the presence of high (2014) and low (2022) sympatric Dorcas gazelle densities revealed very similar estimations (Fig. 5b), suggesting that intra-specific competition had minimal influence on Acacia gazelle dietary considerations.

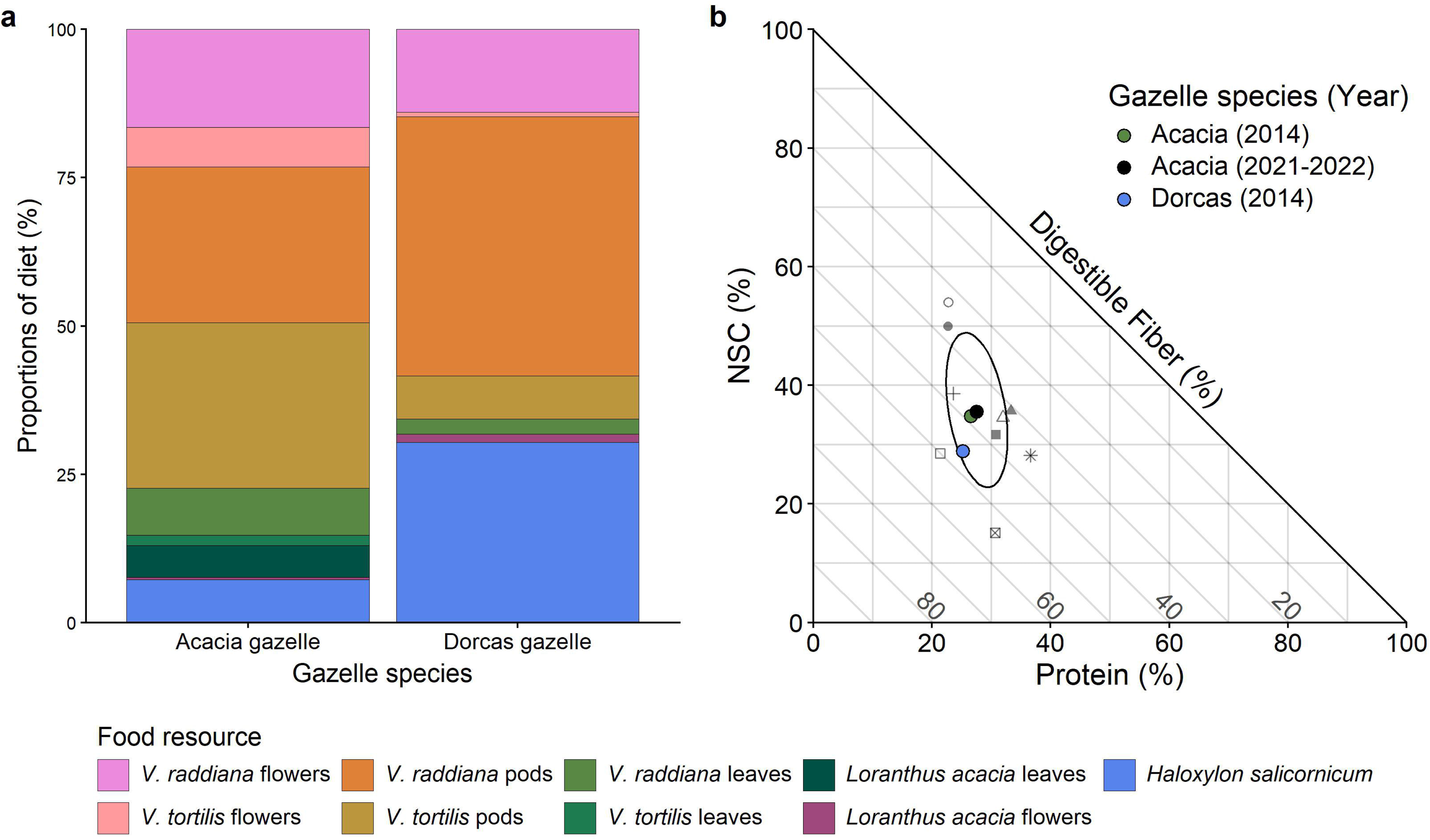

We observed consistent differences in ash and tannin levels between the diets of the two gazelle species. The halophyte *H. salicornicum*, which constituted a major part of Dorcas gazelles’ diet, had higher Na, Mg, and K content (Table S4), hence also significantly higher ash content compared to all other plant resources (*p*<0.03; Table S3). Leaves of the two *Vachellia* trees had significantly higher tannin content compared to all other resources (*p*<0.001 for all; Table S3), suggesting that Acacia gazelles consumed higher quantities of tannins and less salt than Dorcas gazelles.

## 4. Discussion

Our goal was to reveal the dietary considerations of the critically endangered Acacia gazelle to assist population recovery and identify suitable habitats for reintroduction. We found that Acacia gazelles feed almost exclusively on resources from two *Vachellia* trees that differ in their phenology, nutritional composition, and defensive traits. The Acacia gazelles maintained a tight nutritional intake year-round, despite an ever-changing broader nutritional landscape. Specifically, Acacia gazelles regulated their protein intake while selectively foraging for non-structural carbohydrates (NSC). Gazelles preferred browsing on *V. tortilis* individuals with higher protein content, more leaves, and lower density of big prickles over trees with bigger and more fibrous leaves and high density of small prickles. Despite a similar utilization pattern, we failed to identify what dictates the differential use of *V. raddiana* trees. The diet of Acacia gazelles differed substantially from that of the sympatric Dorcas gazelles. While Acacia gazelles used the two *Vachellia* subspecies and fed from their canopies, Dorcas gazelles’ diet included more halophytic shrubs and almost no *V. tortilis* resources. Both gazelles consumed *V. raddiana* resources, but Dorcas gazelles ate primarily from the ground and less from the canopy compared to Acacia gazelles.

Acacia gazelle kept a tight nutritional intake target across seasons despite an ever-changing broader nutritional landscape. Protein intake was about 17% of dry matter consumed across seasons and showed the least variation in comparison to other nutrients. To our best knowledge, such tight regulation of protein consumption in wild herbivores was documented only in primates (Felton et al. 2009b, c, Hou et al. 2021, Takahashi et al. 2021). A common presumption in the herbivore-plant literature is that herbivores must prioritize the consumption of protein-rich foods to fulfill their protein demand (Milton 1979, Mattson 1980, White 1993). This presumption is based on the acknowledged mismatch between the low protein content of many plant resources and the herbivores’ high physiological protein demand. We, however, found no evidence that gazelles attempted to forage selectively for protein to achieve the precise protein intake. Consequently, we suggest that the high protein content across available resources allows these gazelles to maintain a rather stable protein intake while selectively choosing to consume NSCs (Codron et al. 2007, Felton et al. 2016, Ganzhorn et al. 2017).

Acacia gazelles selectively used NSC-rich foods, such as *Vachellia* flowers. Selection of non-protein energy foods has been observed in primates (Rothman et al. 2011) and in other ungulates (Asher et al. 2011, Felton et al. 2016). However, these studies found that attempts to regulate NSC intake come at the expense of protein regulation. We found that Acacia gazelle can simultaneously maintain a similar protein intake while foraging for NSC. To increase NSC intake, gazelles must feed on both *Vachellia* subspecies that provide flowers and ripe fruits at different times. Consequently, a good habitat for Acacia gazelles must include sufficiently large populations of both *Vachellia* subspecies*. V. raddiana* and *V. tortilis* co-occur mainly along the Jordan Rift Valley, south of the Dead Sea (Halevy and Orshan 1972). This may explain why the historic distribution of Acacia gazelles was limited to this area (Mendelssohn 1974), while the distribution of the sympatric Dorcas gazelle is much wider and includes diverse desert habitats (Yom-Tov 2016).

When examining diets and food preferences researchers often use resource species as a basic unit, neglecting intraspecific variability that could be substantial (Lawler et al. 1998, Chapman et al. 2003, Rothman et al. 2012, Wam et al. 2018). Acacia gazelles chose to forage on specific trees and ignored others, likely due to intraspecific variation in tree traits. The preferred *V. tortilis* trees were richer in protein and NSC relative to fibers and had high densities of small leaves and lower densities of prickles relative to the un-browsed trees. We hypothesized that in their attempt to improve foraging efficiency, gazelles consider both the nutritional intake and the feeding costs. Understanding palatability variation across trees is necessary when estimating an area’s carrying capacity since not all potential resources are being used in similar ways (DeGabriel et al. 2014, Youngentob et al. 2026). Moreover, the nutritional content and chemical and physical defenses can vary with tree age and in response to browsing pressure (Gowda 1997, Rohner and Ward 1997, Fornara and Du Toit 2007, Nzimande et al. 2022). Thus, gazelles may experience nutritional limitations even if the number of trees and the foliage availability may erroneously suggest ample forage. We suggest that the nutritional and defensive traits of trees should be monitored to prevent such convoluted density-dependent reduction in resource quality and accessibility. This recommendation is especially relevant when the population is fenced.

The diet of the sympatric Acacia and Dorcas gazelles varied substantially, suggesting limited interspecific competition for food resources. Our continuous observation of Acacia gazelle individual-foraging from the summer of 2014 revealed a similar diet composition and nutritional intake to that observed in the summer of 2022 using a different observation approach. We found that Acacia gazelles consumed foliage, flowers, and pods from the canopy of the two *Vachellia* subspecies, and ate the abundant parasitic plant *Loranthus acaciae*. In comparison, the diet of the sympatric Dorcas gazelle included almost exclusively *V. raddiana* resources collected from the ground. Halophytic shrubs constituted 30% of the Dorcas gazelle’s diet in comparison to only 7.2% in the Acacia gazelle’s diet. Despite these large dietary differences, their protein intake converged to similar values, and their intake of carbohydrate sources differed only slightly between the two gazelle species, suggesting similar physiological needs.

Similar physiological needs of sympatric species that feed on different diets may suggest dietary niche separation that requires specific adaptations. Acacia gazelles forage high in the canopy by standing only on the hind legs or by leaning with the front legs on low branches (Blank 2005). These gazelles have to cope with high tannin levels and presumably other chemical defenses and avoid the spines and prickles, especially those of *V. tortilis*. On the other hand, Dorcas gazelles need to cope with high salt levels and fluctuations in food availability. Early studies on Dorcas gazelle physiology support these insights. As a desert specialist, this species can double its urine concentration (Ghobrial 1974) and maintain body weight while drinking water with up to 2% salinity (Ghobrial 1976). Our results suggest that the two sympatric gazelle species evolved to consume different diets, hence experience low interspecific competition. Our findings that Acacia gazelles’ diet and nutritional intake did not differ before and after the sharp drop in the sympatric Dorcas gazelle population density coincide with this conclusion.

Our comparative observations were conducted during summer and fall. Thus, our inferences may be limited to these seasons. We also found that Acacia gazelles increased consumption of *V. raddiana* in the winter, possibly intensifying competition with the Dorcas gazelles. However, previous work showed that Dorcas gazelles tend to substantially decrease the use of *V. raddiana* during the winter (Baharav 1980, 1982), supporting our conclusion of dietary niche separation between the two gazelle species. Acacia gazelles specialize in feeding from *Vachellia* trees and switch between subspecies and their resources to maintain a tight nutritional target. Tracking both *Vachellia*s’ phenology may allow this gazelle species to avoid seasonal nutritional shortages in this unpredictable hyper-arid environment. Moreover, *Vachellia* trees supply resources that are less prone to fluctuations in availability between years compared to annual plants or small perennial shrubs. Stability in resource quality and quantity may support a high reproductive rate allowing Acacia gazelles to calve in all seasons and even twice a year (Shalmon 1991, Mendelssohn et al. 1997, Blank 2000). Other gazelle species, including the sympatric Dorcas gazelle, exhibit a synchronized seasonal reproduction peak that is tightly timed to match the annual peak in resource availability (Baharav 1983, Yom-Tov 2016). Thus, Acacia gazelles may have greater reproductive potential than Dorcas gazelles, but, are limited to a more specific habitat where the two *Vachellia* subspecies are present.

### 4.1 Conclusions and management implications

We used a nutritional geometry approach to uncover Acacia gazelles’ dietary considerations in attempts to guide conservation efforts. Acacia gazelles forage predominantly on two *Vachellia* subspecies that provide crucial resources at different seasons. Acacia gazelle’s macro-nutritional intake varied little between months in 2021-2022 and was similar to that observed in the summer of 2014. This enforces confidence in the estimated intake representing nutritional needs and suggests that, since Acacia gazelles were able to reach it year-round, currently, the population does not suffer from a nutritional deficiency that requires supplemental feeding. However, in case of future needs, identifying this precise intake target may enable managers to provide feed that meets the species-specific needs. Acacia gazelles’ specialization on *Vachellia* resources poses a potential obstacle to the population’s recovery, since the two *Vachellia* subspecies coexist only in alluvial fans along the Jordan Rift Valley, south of the Dead Sea. This restricted habitat is diminishing in an unprecedented way due to agricultural expansion. Reintroductions of Acacia gazelles may thus necessitate a broader conservation plan to expand the protection of their unique habitat, and restore the nutritional landscape that can support their populations (Youngentob et al. 2026). It was hypothesized that similar physiological needs may induce intense interspecific competition for food between the two sympatric gazelles (Shalmon et al., 2020). Instead, our findings reveal low interspecific competition between the two gazelles. Thus, we recommend postponing the seemingly unjustified removal of Dorcas gazelles from the enclosure that may counterintuitively harm the Acacia gazelles by denying potential anti-predatory benefits (e.g., social vigilance, dilution effect). We also found that Acacia gazelles selectively feed on certain *Vachellia* individuals and identified the possible traits that affect this selection for *V. tortilis* but not for *V. raddiana*. Since trees subject to different browsing pressures may change the expression of these traits, continuous monitoring is important. We recommend monitoring *Vachellia* recruitment, canopy height, and defensive traits to prevent density-dependent limitations and identify otherwise obscured reductions in resource accessibility and quality. Developing state-of-the-art hyperspectral remote sensing tools to assess the nutritional landscape over an extended spatial scale would be an important step in this direction. Our case study highlights the benefit of adopting a nutritional ecology approach to guide management and reintroduction plans for endangered wildlife.

