## Supplementary material for "A recipe for recovery: Diet as a decisive tool in species conservation": Fig. S; Table S; Appendix

### **Supplementary materials**

**Appendix 1 - Observations protocol:**

We used two observation points during the study, following the Acacia gazelles' daily activity pattern. To minimize our disturbance, we conducted morning observations from a hide within the enclosure, to which we entered before first light, and exited only after the gazelles left the area. To avoid scaring off the gazelles in the afternoon, we avoided entering the enclosure and observed the gazelles from a hill that overlooks the entire northern part of the enclosure. We entered the hides 30-60 minutes before the gazelles' foraging activity rumped up to further minimize potential disturbance. The morning sessions started before the first light and ended when the herd left the northern area. We began the evening sessions before the gazelles began moving toward the northern part of the enclosure, and finished the observation after dark. During each observation, we observed gazelle females and calves using a telescope (82mm x20-60) and a binocular (10x42). In 2021-2022, we conducted 58 group observation sessions for a total of 123 hours and 25 minutes, with a mean session of 128±42 mins. Group size varied between 1 to 12 individuals. In 2014 summer, we collected 76 individual focal observations of 15±9 mins, of which were 38 adult female Acacia gazelles, 19 young Acacia gazelles, and 19 adult female Dorcas gazelles. Each tree or large bush observed in 2021-2022 in the northern part of the enclosure received a unique record number with detailed information including species, exact location, physical characteristics, and a photograph. During an observation, we identified the individual plants eaten on an aerial photo by pinning their location based on measured azimuth and distance, using a mobile QField application, a laser rangefinder (Nikon Monarch 3000 Stabilized), and a compass. We extracted all locations and their feeding data into a QGIS project. At the end of each morning session, we approached the consumed plants, verified our location identification, and collected samples from all the plants that were consumed that morning or during the evening observation of the preceding day.

**Table S1 Amount and kind of plant samples collected in the northern part of the gazelles' enclosure.**

| **Species** | **Part** | **Total number of samples** | **Analyzed by wet chemistry** |
| --- | --- | --- | --- |
| *V. raddiana* | Leaves | 97 (16)^[1]^ | 40 (8)^[2]^ |
| *V. raddiana* | Pods | 89 (3)^1^ | 60 (15)^2^ |
| *V. raddiana* | Flowers | 21 (4)^1^ | 8 |
| *V. tortilis* | Leaves | 72 (19)^1^ | 30 (9)^2^ |
| *V. tortilis* | Pods | 23 | 21 (5)^2^ |
| *V. tortilis* | Flowers | 15 | 8 |
| *Loranthus acacia* | Leaves | 10 (3)^1^ | 10 |
| *Loranthus acacia* | Flowers | 17 | 8 |
| *Loranthus acacia* | Fruits | 7 | 7 |
| *Haloxylon salicornicum* | Leaves | 9 | 9 |
| *Lycium shawii* | Leaves | 8 | 4 |
| *Salsola imbricata* | Leaves | 3 | 1 |
| *Nitraria retusa* | Leaves | 2 | 2 |
| *Tamarix sp.* | Leaves | 1 | 1 |
| **Total** |  | **400** | **209** |

**Appendix 2 - NIRS models (leaves and pods of *Vachellia* subspecies):**

For the macro-nutrients models, we centered and standardized the spectra and applied a Standard Normal Variate correction (SNV) to account for potential differences in particle size. To reveal hidden peaks in the spectra we used the second derivative with a Norris derivative filter using a segment length of 7 points and gaps of 5 points between segments. For the tannins model, we used a constant path length, which produced better results than the SNV correction, and used the smoothed second derivative with a segment length of 5 and gap of 4. We used a PLS regression to create the predictive model, this method is preferable since it considers both the standards' spectra and evaluated components, and is known to produce good calibration results (Agelet & Hurburgh 2010). For each component, we used the minimal number of latent variables (factors) that minimized the predicted residual error sum of squares (PRESS) produced by "leave-one-out" cross-validation. We evaluated the model performance by considering the root mean square error of the calibration (RMSEC), cross-validation (RMSECV), and external validation set (RMSEP), see Table S1 for the number of samples used. We chose the model that minimized errors and the differences between the calibration and predicted errors of the cross and external validation to avoid overfitting. Models’ performances are presented in Table S2.

**Table S2. Statistical measures and performance of macro-nutrients and tannins NIRS models.**

|  | Protein | NDF | ADF | ADL | Lipids | NSC | Tannins |
| --- | --- | --- | --- | --- | --- | --- | --- |
| RMSEC^[3]^ | 0.90 | 2.35 | 1.90 | 1.96 | 0.58 | 2.93 | 0.18 |
| *R^2^* | 0.96 | 0.96 | 0.96 | 0.87 | 0.94 | 0.89 | 0.94 |
| RMSEferP^1^ | 1.26 | 2.41 | 2.18 | 1.88 | 0.71 | 3.93 | 0.19 |
| *R^2^_P_* | 0.92 | 0.96 | 0.95 | 0.86 | 0.92 | 0.80 | 0.95 |
| RMSECV^1^ | 1.64 | 3.21 | 2.94 | 2.46 | 0.69 | 4.33 | 0.22 |
| *R^2^_CV_* | 0.86 | 0.93 | 0.90 | 0.78 | 0.92 | 0.75 | 0.91 |
| Number of factors | 7 | 5 | 4 | 4 | 5 | 5 | 4 |

**Supplementary tables:**

**Table S3. Mean ± SD (% of dry matter, tannins were measured as absorbance values) of the food resources and intake of Acacia gazelles. *Vachellia*s leaves and pods values were obtained from NIRS predictive models for NSC, lipids, NDF, ADF, ADL, and tannins (Table S2), ash content was quantified by wet chemistry only. All other resources were quantified by wet chemistry. Hemicellulose and cellulose were calculated as NDF-ADF and ADF-ADL, respectively.**

| **Species** | **Part** | **Crude Protein** | **NSC** | **Lipids** | **NDF** | **Hemicellul-ose** | **ADF** | **Cellulose** | **ADL (Lignin)** | **Ash** | **Tannins** |
| --- | --- | --- | --- | --- | --- | --- | --- | --- | --- | --- | --- |
| ***V. raddiana*** | **leaves** | 16.0±1.85**^a^** | 20.4±5.83**^bc^** | 4.34±0.979**^c^** | 34.9±4.95**^bc^** | 8.87±3.33**^b^** | 26.0±3.57**^bc^** | 10.6±1.48**^c^** | 15.4±3.05**^c^** | 8.97±2.54**^a^** | 1.3±0.44**^c^** |
| ***V. tortilis*** | **leaves** | 18.0±1.81**^a^** | 22.8±5.49**^b^** | 5.14±1.15**^d^** | 31.8±4.91**^bd^** | 7.68±2.66**^bc^** | 24.1±4.19**^bcd^** | 9.85±1.83**^c^** | 14.3±3.24**^c^** | 8.32±2.37**^a^** | 1.1±0.49**^c^** |
| ***V. raddiana*** | **pods** | 17.1±3.25**^a^** | 17.3±4.02**^a^** | 1.57±0.784**^a^** | 44.0±6.42**^a^** | 10.3±1.66**^a^** | 33.7±5.18**^a^** | 20.4±3.52**^a^** | 9.93±2.17**^a^** | 7.71±3.08**^a^** | 0.49±0.32**^a^** |
| ***V. tortilis*** | **pods** | 21.6±2.17**^b^** | 21.9±4.34**^bc^** | 2.69±0.523**^b^** | 33.7±4.91**^bc^** | 8.15±1.62**^b^** | 25.5±2.70**^b^** | 17.3±1.47**^b^** | 8.11±1.83**^b^** | 6.94±1.95**^a^** | 0.23±0.10**^b^** |
| *V. raddiana* | flowers | 14.5±2.30 | 34.6±6.51 | 3.98±0.531 | 27.3±3.21 | 5.68±1.69 | 21.6±2.20 | 9.18±1.67 | 12.5±1.82 | 7.31±2.18 | 0.57±0.20 |
| *V. tortilis* | flowers | 15.1±2.01 | 33.4±8.62 | 3.71±0.669 | 29.3±5.49 | 4.68±5.91 | 24.6±9.60 | 13.6±8.59 | 11.0±1.55 | 8.84±2.63 | 0.43±0.14 |
| ***L. acacia*** | **leaves** | 11.7±5.48**^c^** | 15.8±3.58**^acd^** | 3.67±1.02**^cd^** | 22.9±4.93**^d^** | 3.69±1.93**^c^** | 19.2±4.35**^d^** | 10.2±1.37**^cd^** | 9.02±3.42**^ab^** | 8.24±2.12**^a^** | 0.41±0.083**^a^** |
| *L. acacia* | fruits | 9.05±1.38 | 26.8±4.97 | 16.5±3.18 | 19.7±2.85 | 6.76±1.20 | 12.9±1.85 | 7.36±1.07 | 5.56±0.876 | 5.93±4.26 | 0.29±0.072 |
| *L. acacia* | flowers | 11.2±1.47 | 17.9±3.01 | 5.64±1.41 | 26.2±1.61 | 8.03±1.34 | 18.2±1.40 | 9.13±1.07 | 9.03±1.05 | 7.19±1.30 | 0.13±0.019 |
| ***H. salicornicum*** | **leaves** | 18.8±2.34**^ab^** | 9.66±0.782**^d^** | 2.30±0.709**^ab^** | 37.3±3.69**^ac^** | 18.5±1.49**^d^** | 18.9±2.96**^cd^** | 13.8±2.23**^d^** | 5.07±0.785**^b^** | 17.5±2.47**^b^** | 0.00±0.00**^d^** |
| *L. shawii* | leaves | 24.9±7.85 | 15.8±3.97 | 3.84±0.909 | 29.2±3.18 | 10.6±0.951 | 18.6±3.23 | 8.42±1.86 | 10.2±3.92 | 24.1±7.17 | 0.00±0.00 |
| *S. imbricata* | leaves | 26.6 | 8.03 | 2.38 | 31.0 | 18.2 | 12.8 | 8.15 | 4.65 | 19.0 | 0.01 |
| **Intake** ^[4]^ | | 17.0±1.51 | 21.9±4.07 | 3.69±0.95 | 35.0±4.03 | 8.96±2.56 | 26.0±2.82 | 13.6±5.76 | 12.2±2.22 | 9.76±3.82 | - |
| **CV** ^[5]^ | | 0.089 | 0.19 | 0.26 | 0.12 | 0.29 | 0.12 | 0.42 | 0.18 | 0.39 | - |

**Table S4. Mineral content (% dry matter) of food resources consumed by both gazelle species in 2014. Letters are Conover test results between food resources observed for each mineral.**

| **Species** | **Part** | **Ca** | **P** | **Na** | **Mg** | **K** |
| --- | --- | --- | --- | --- | --- | --- |
| *V. raddiana* | leaves | 2.06±0.83**^a^** | 0.126±0.0232**^ac^** | 0.0863±0.0463**^a^** | 0.261±0.120**^ab^** | 0.72±0.178**^b^** |
| *V. tortilis* | leaves | 1.98±0.74**^ab^** | 0.148±0.0402**^ab^** | 0.0801±0.0646**^a^** | 0.304±0.0585**^b^** | 0.76±0.199**^b^** |
| *V. raddiana* | pods | 1.18±0.55**^bc^** | 0.193±0.0517**^b^** | 0.0489±0.0474**^ab^** | 0.241±0.0431**^ab^** | 1.19±0.398**^c^** |
| *V. tortilis* | pods | 0.92±0.23**^c^** | 0.301±0.0322**^d^** | 0.0169±0.00440**^b^** | 0.292±0.0444**^b^** | 1.28±0.103**^c^** |
| *L. acacia* | leaves | 1.23±0.23**^abc^** | 0.140±0.0458**^abc^** | 0.122±0.239**^ab^** | 0.185±0.0598**^a^** | 2.04±0.566**^a^** |
| *H. salicornicum* | leaves | 1.43±0.29**^abc^** | 0.0988±0.0223**^c^** | 1.67±0.321**^c^** | 1.11±0.112**^c^** | 2.50±0.441**^a^** |

**Table S5. Results of *V. tortilis* PLS-DA variable importance permutations showing the mean Mahalanobis (*M_d_*), the fraction of permuted models with significant cluster separation, and means of fit measures (*R^2^_y_*, *DR^2^_y_*). Variables with measures smaller than the original (Unpermuted) model are highlighted, original model's measures are presented for comparison.**

| Variable permuted | Mean *M_d_* | Fraction of significant models | Mean *R^2^_y_* | Mean *DR^2^_y_* |
| --- | --- | --- | --- | --- |
| **Hemicellulose** | **2.36** | **0.023** | **0.60** | **0.63** |
| **Big prickles** | **2.87** | **0.27** | **0.69** | **0.72** |
| **Protein** | **2.91** | **0.35** | **0.70** | **0.73** |
| Lignin | 3.06 | 0.75 | 0.72 | 0.75 |
| Lipids | 3.30 | 1 | 0.75 | 0.78 |
| Small prickles | 3.38 | 1 | 0.76 | 0.79 |
| Leaf width | 3.38 | 1 | 0.76 | 0.80 |
| Leaf length | 3.45 | 1 | 0.77 | 0.81 |
| Cellulose | 3.47 | 1 | 0.77 | 0.80 |
| Leaves | 3.49 | 1 | 0.77 | 0.81 |
| Tannins | 3.55 | 1 | 0.78 | 0.81 |
| NSC | 3.56 | 1 | 0.78 | 0.81 |
| Big spines | 3.61 | 1 | 0.78 | 0.81 |
| Small spines | 3.62 | 1 | 0.78 | 0.82 |
| **Unpermuted model** | **3.28** |  | **0.75** | **0.78** |

**Supplementary figures:**

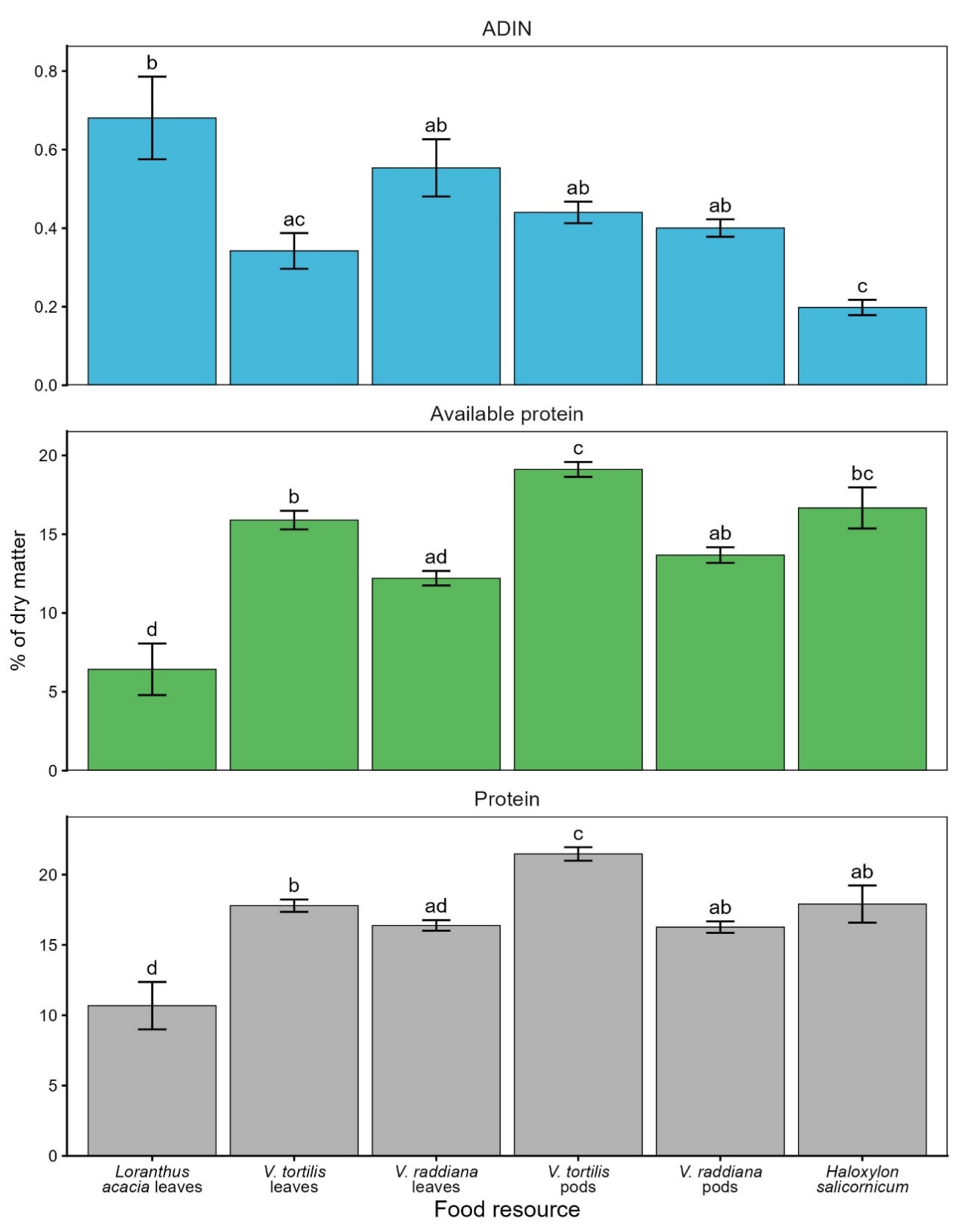

Fig. S1 Results of Acid Detergent Insoluble Nitrogen (ADIN), available protein (calculated by multiplying the ADIN by the 6.25 conversion factor and subtracting from the crude protein measure), and crude protein contents of major food resources. Letters are the Conover test results between food resources observed for each measure. The analyses were done on 139 samples, as we were unable to calibrate an acceptable NIRS model of ADIN. ADIN concentrations did not differ between *Vachellia* resources. Also, the relative differences between resources did not change when comparing the available protein to crude protein, aside from *H. salicornicum* being significantly higher in available protein than *V. raddiana* leaves and pods.

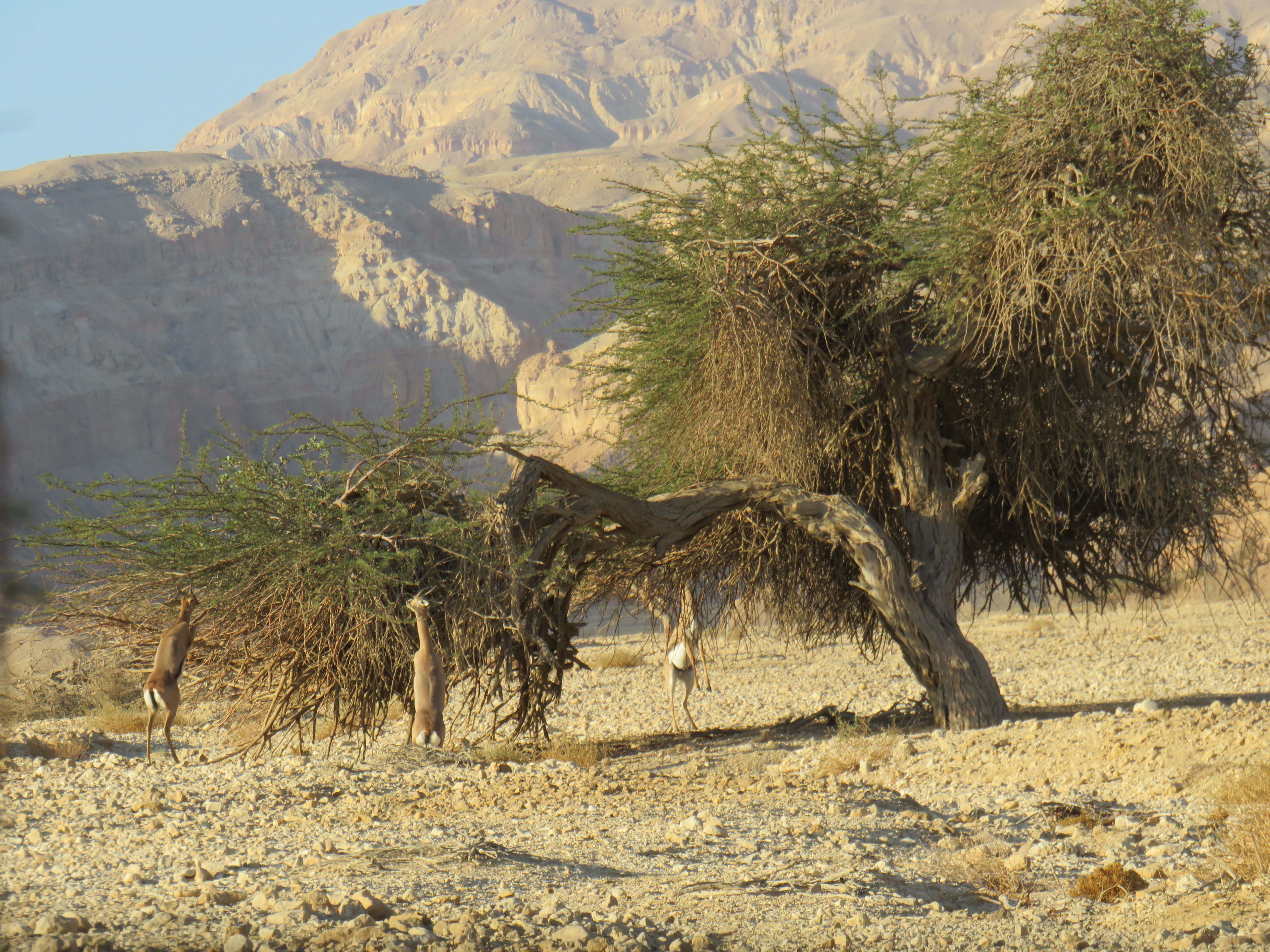

**Fig. S2 Adult female (center) and males (adult on the right and young on the left) Acacia gazelles standing on their hind legs (bipedal) while feeding. Max feeding height was estimated in the field.**

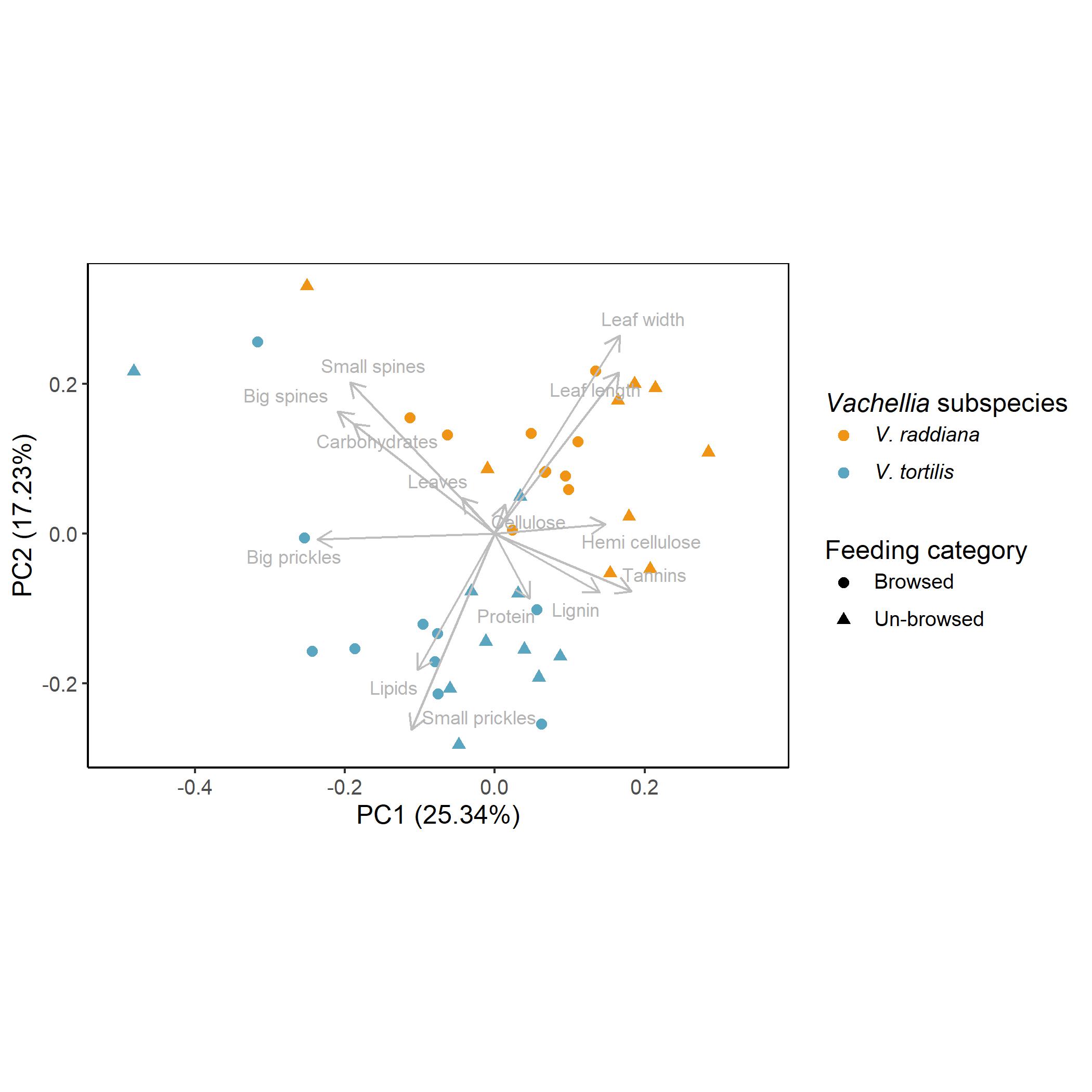

**Fig. S3 PCA ordination biplot showing chemical and physical differences between *Vachellia* subspecies.**

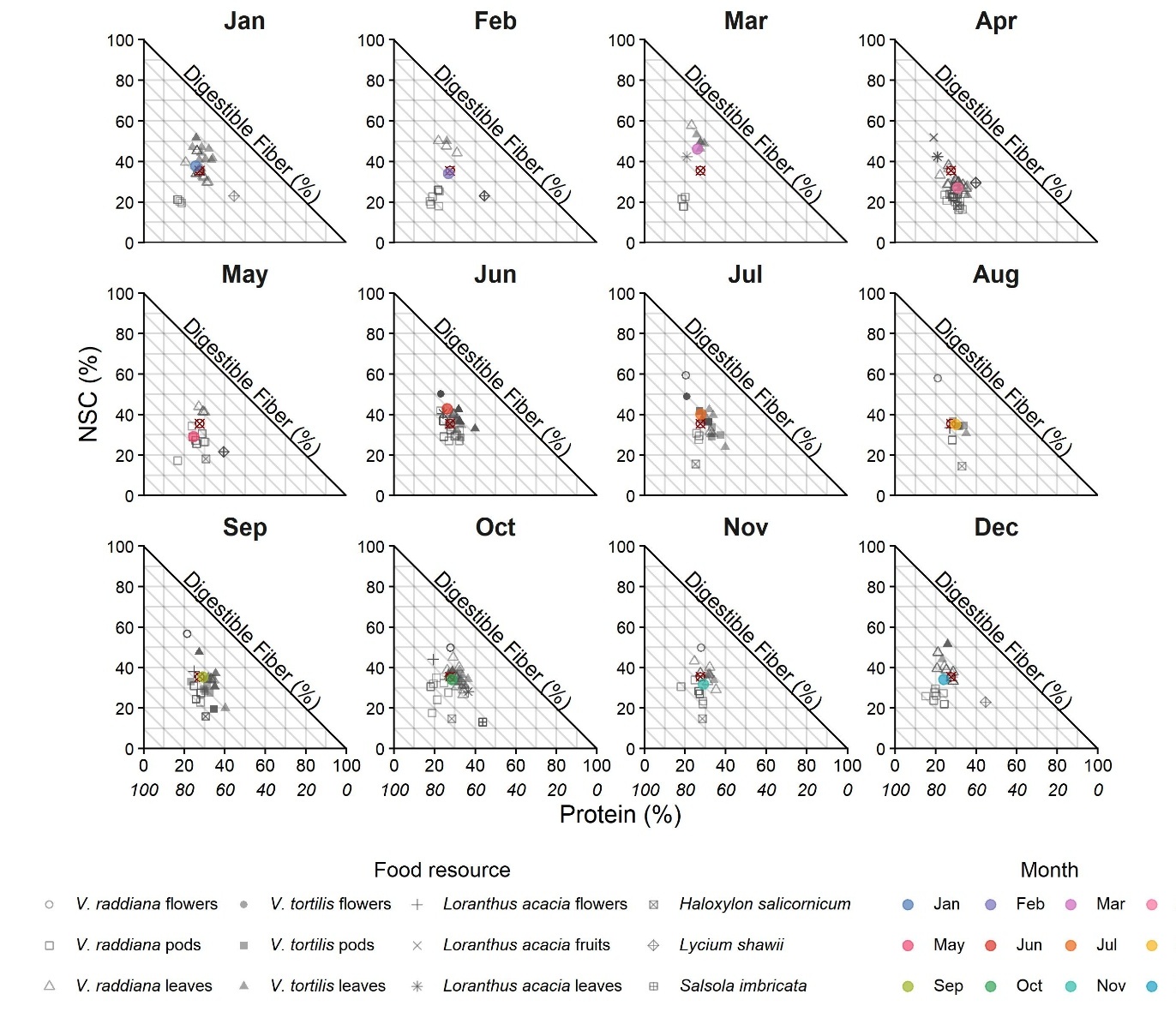

**Fig. S4 Acacia gazelle's monthly nutritional intake and resources. Red symbol represents the annual (mean) nutritional target and points colored by month are the monthly mean nutritional intake. In gray are all the different resources collected during that month. The top X-axis is the protein horizontal axis, and the bottom italic axis is the Z (Digestible fiber) diagonal axis.**

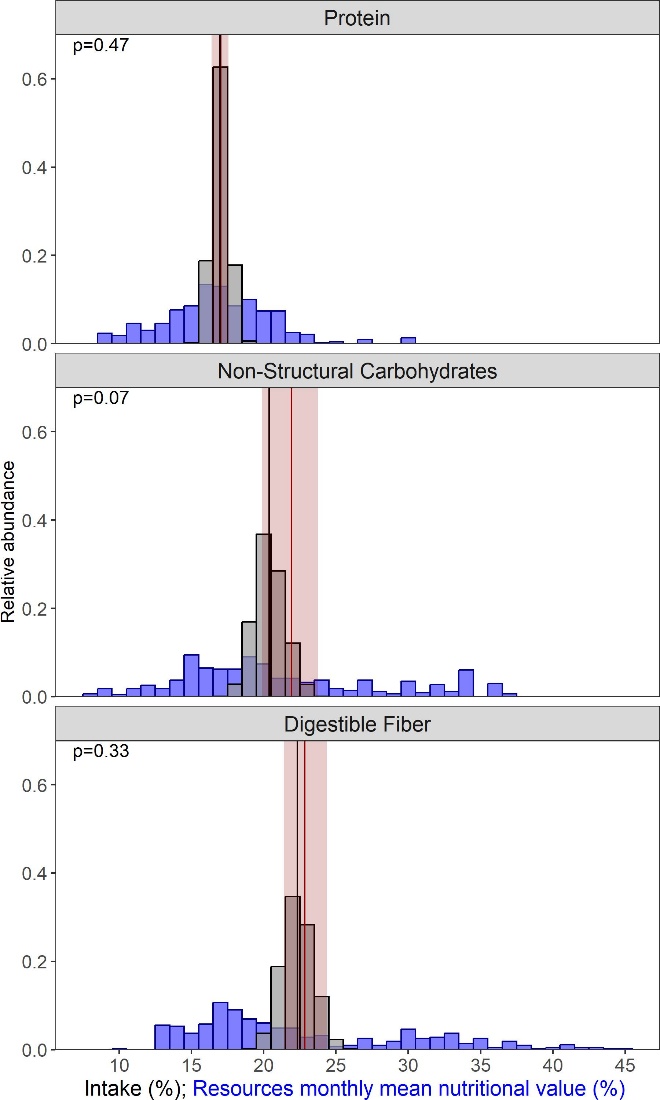

**Fig. S5 Results of the permutation test of gazelles’ selection for a macro-nutrient. Selection for a macro-nutrient was assessed as whether the gazelle observed consumption deviates from random feeding. The distributions of Acacia gazelles' random intakes as obtained with 1000 permutations (gray), with the null distribution median marked in a black line. The observed intake target is marked with a red line, and the shaded area is the 95% bootstrap confidence interval, estimated by resampling observation sessions. The distribution of mean monthly nutritional values of food resources is displayed in blue. P-value was calculated as the fraction of permuted values exceeding the observed intake, and is reported as the median across 1000 permutation runs. Digestible fiber are calculated as NDF-ADL and include hemicellulose and cellulose.**

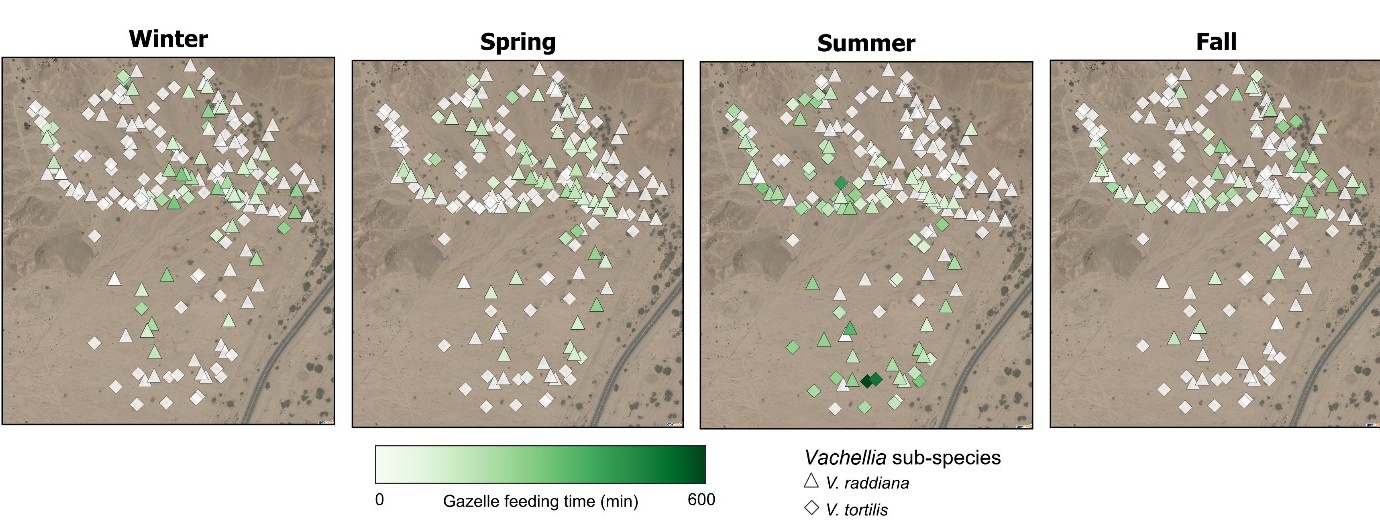

**Fig. S6 *Vachellia* trees spatial distribution and utilization by Acacia gazelles. Each symbol is a single mapped tree (triangles for *V. raddiana* and diamonds for *V. tortilis*). Trees are colored by the total time the gazelles spent feeding from it in each season - winter (December-February), spring (March-May), summer (June-September), and fall (October-November).**

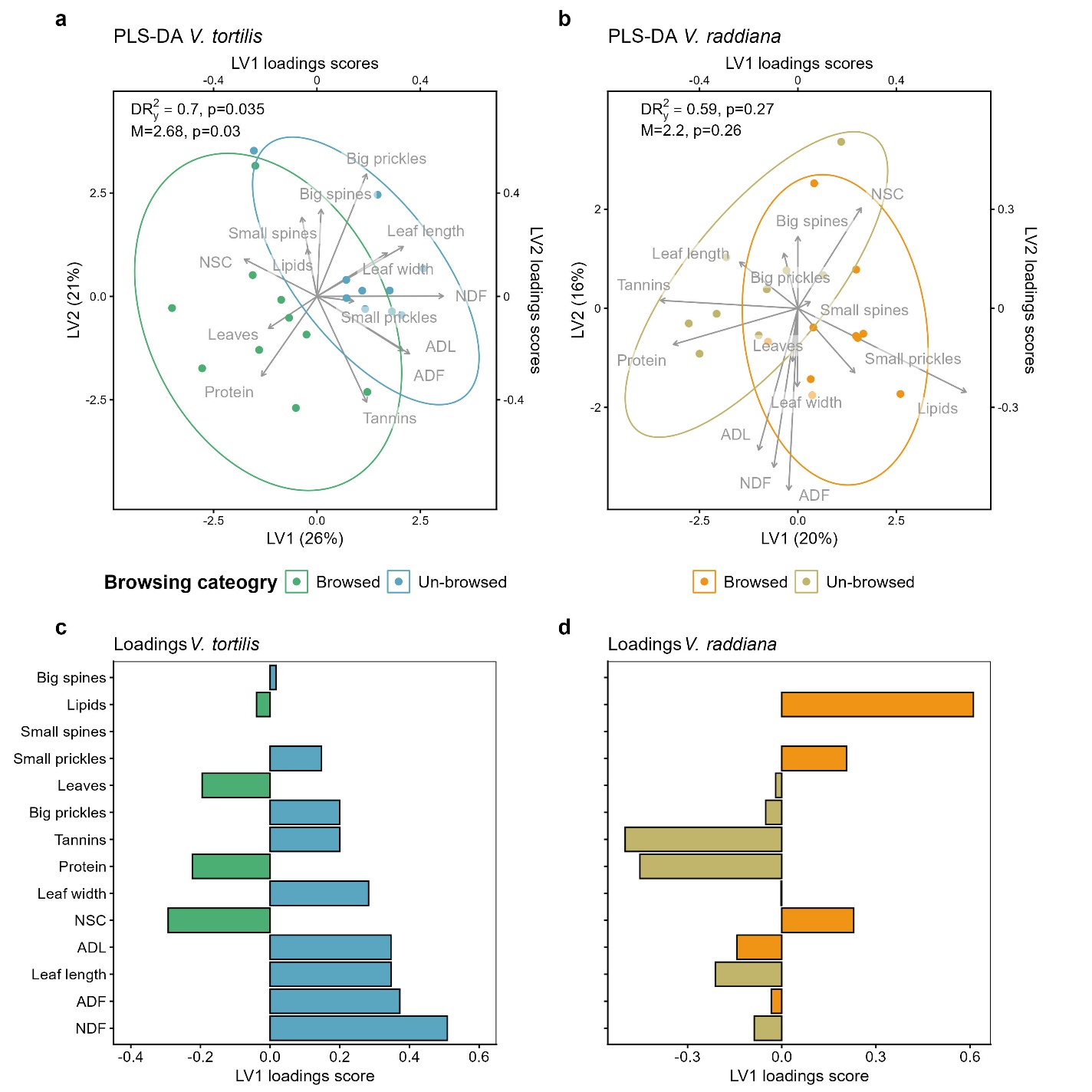

**Fig. S7 PLS-DA ordination results for the analytical fiber fractions. a;b. biplots for both *Vachellia* subspecies. Gray arrows' sizes are equivalent to the loading scores for each ordination variable. Percentages on the axes are the variation explained by each. c;d. Loading scores of the first component colored by the browsing category in which the median is maximum for each variable.**

1. Numbers in parenthesis are the subset of samples collected from trees that had no feeding observations.
2. Numbers in parenthesis are the subset of samples that were used for external validation of NIRS model. The rest were used for calibration.
3. $RMSE=\frac{\sqrt{\sum_{p=1}^{p} \sum_{n=1}^{n} \left( \hat{y}_{n,p}-y_{n,p} \right)^{2}}}{n\cdot p}$

   Where *p* is the number of factors used for each component and *n* is the number of samples. $\hat{y}$ is the predicted value of the component by each factor and *y* is the true value.
4. Letters are significant Conover test results between food resources observed in 2014 (in bold) for each macro-nutrient.

   Mean intake was weighted for each day by the time spent feeding and then averaged across days. Intake was not calculated for tannins as the results are qualitative
5. coefficient of variance - calculated as SD/mean
